# Quinoa as Functional Food? Urinary elimination of ecdysterone after consumption of quinoa alone and in combination with spinach

**DOI:** 10.1101/2023.10.10.560427

**Authors:** Eduard Isenmann, Tasha Yuliandra, Konstantina Touvleliou, Matthias Broekmann, Xavier de la Torre, Francesco Botrè, Patrick Diel, Maria Kristina Parr

## Abstract

The phytosteroid ecdysterone is included on the monitoring list of the World Anti-Doping Agency. Therefore, the consumption of food rich in ecdysterone is the focus of a lively debate. Thus, urinary excretion of ecdysterone and its metabolites in humans was investigated following quinoa consumption alone and in combination with spinach.

After intake of both preparations, ecdysterone and two metabolites were excreted in urine. Maximum concentrations of ecdysterone ranged from 0.44–5.50µg/mL after quinoa and 0.34– 4.09µg/mL after quinoa with spinach. The total urinary excreted amount as parent drug plus metabolites was 2.60(1.09)% following quinoa and 1.71(0.86)% after combination. Significant differences were found in total urinary excreted amounts of ecdysterone, 14-deoxy-ecdysterone, and 14-deoxy-poststerone. In conclusion, only small proportions of ecdysterone from quinoa and the combination with spinach were excreted in urine. The results indicate that both, quinoa and spinach, are poor sources of ecdysterone.

## 1. Introduction

Since 2020, ecdysterone, an ecdysteroid hormone, is included in the monitoring program of the World Anti-Doping Agency classified as an anabolic agent (World Anti-Doping Agency). Its discovery was traced back to the 1950s in insects; years later, it was found in plants in higher amounts (Bathori et al., 2008; Butenandt & Karlson, 1954). Various pharmacological activities of ecdysterone were reported since then, for instance, its anabolic activity has been investigated by *in-silico*, *in-vitro*, and *in-vivo* studies (Gorelick-Feldman et al., 2008; Isenmann et al., 2019; Parr et al., 2015; Parr et al., 2014; Syrov, 2000; Toth et al., 2008). *In-silico* and *in-vitro* studies reported that the anabolic activity of ecdysterone is mediated by estrogen receptor beta (ER-β), thus no androgen-related side effect was observed (Parr et al., 2014). Additionally, *in-vitro* studies in mouse and human skeletal muscle cells showed a significant increase in protein synthesis after treatment with ecdysterone (Gorelick-Feldman et al., 2008; Parr et al., 2014). An increase in muscle fiber size, body weight, protein content, and enhancement in muscle strength was also observed in rats after administration of ecdysterone (Gorelick-Feldman et al., 2008; Parr et al., 2015; Parr et al., 2014; Syrov, 2000; Toth et al., 2008). Furthermore, long-term supplementation of ecdysterone showed an increase in muscle mass and performance enhancement during resistance training in healthy athletes (Isenmann et al., 2019).

In anti-doping laboratories, the use of urine as biological samples is common, easy to collect, efficient, and non-invasive compared to serum samples (Nicoli et al., 2016). The urine samples are used to detect a compound or its metabolites that is prohibited in sports. The presence of the compounds and the metabolites in urine can be used as an indicator of doping usage. Several pharmacokinetic studies of ecdysterone have been conducted in humans, especially those related to urinary excretion (Ambrosio et al., 2021; Brandt, 2003; Dioh et al., 2023; Parr et al., 2019; Tsitsimpikou et al., 2001). After oral administration of ecdysterone, the parent compound and metabolites were excreted in urine such as 14-deoxy-ecdysterone, 2-deoxy-ecdysterone and deoxy-ecdysone (Ambrosio et al., 2021; Dioh et al., 2023; Tsitsimpikou et al., 2001). This is in line with own observations (Ambrosio et al., 2021; Parr et al., 2019). Therefore, both ecdysterone and these reported metabolites may be used as markers for the detection of ecdysterone consumption.

However as mentioned earlier, ecdysterone is also present in edible plants consumed in a normal diet such as quinoa *(Chenopodium quinoa)* and spinach *(Spinacia oleracea)*. In quinoa seeds, the amount of ecdysterone varies between 138 – 804 µg/g seed (Graf et al., 2014; Graf et al., 2016; Kumpun et al., 2011). The variation of ecdysterone content is reportedly related to seed variety, genetic, and environmental conditions (Graf et al., 2016; Kumpun et al., 2011). Furthermore Graf et al., reported that there is a positive correlation between oil content and ecdysterone content in quinoa seeds (Graf et al., 2016). Together with ecdysterone as the main ecdysteroid, Zhu et al. and Kumpun et al. were able to isolate different makisterone A metabolites such as 24-*epi*-makisterone A, 24(28)-dehydromakisterone A, and 20,26-dihydroecdysone (Zhu et al., 2001), 24,25-dehydroinokosterone, 25,27-dehydroinokosterone, and 5β-hydroxy-24(28)-dehydromakisterone A from quinoa (Kumpun et al., 2011). In addition, Nsimba et al. found 20,26-dihydroxy,28-methyl ecdysone, 20,26-dihydroxy,24(28)-dihydroecdysone, and 20-hydroxyecdysone 22-glycolate (Nsimba et al., 2008). Dini et al. investigated the presence of ecdysteroids in Kancolla seeds, a sweet variety of quinoa, and isolated a new compound, kancollosterone (Dini et al., 2005).

Based on high concentrations of ecdysterone in quinoa, effects against obesity in mice were observed (Foucault et al., 2012; Graf et al., 2014; Noratto et al., 2019). Initial studies with elderly humans found similar effects on body weight, BMI, LDL, and total cholesterol concentrations (Pourshahidi et al., 2020). In addition, the consumption of 25 g of quinoa flakes for 4 weeks in overweight postmenopausal women showed a decrease in total cholesterol, serum triglycerides, and LDL levels (De Carvalho et al., 2014). Ecdysteroids from quinoa also have antioxidant activity and can be used in the treatment of skin and ageing (Graf et al., 2015; Nsimba et al., 2008).

Similar to quinoa, the amount of ecdysteroids in field-grown spinach leaves, is ranging from 4 to 230 µg/g fresh weight (FW) and in laboratory-grown plants up to 800 µg/g FW (Grebenok & Adler, 1991). Furthermore, in addition to ecdysterone, spinach contains related compounds such as polypodin B, makisterone A, 2-deoxy-ecdysterone, and a small amount of ecdysone (Bakrim et al., 2008; Fang et al., 2022; Grebenok & Adler, 1991, 1993).

Different seeds of spinach from several locations grown under controlled conditions showed the amount of ecdysterone from the leaves varies between 9.3 – 17.3 µg/g dry weight (DW) (Gorelick et al., 2020). Fresh, cooked, and frozen leaves of spinach also revealed a significant variation in the content of ecdysterone, 80 to 428 µg/g dry mass reported by Fang et al. (Fang et al., 2022), 17 – 885 µg/g dry mass based on the study of Bajkacz (Bajkacz et al., 2020), and 24 µg/g as investigated by Chen et al. (Chen & Feng, 2021). On the other hand, only a small amount of ecdysterone (0.1 µg/g FW) was quantified from fresh spinach as reported by Grucza et al. (Grucza et al., 2021). The high variation in the content of ecdysterone in spinach is related to the location of the growing plants, the genetic information, the growing season, the development stage of plants, and plant variety (Bakrim et al., 2008; Gorelick et al., 2020; Grebenok & Adler, 1991).

Spinach also contains various phytochemicals that are beneficial for human health such as flavonoids, carotenoids, fatty acids, multivitamins, and minerals. Because of its contents, spinach has been proven to have antioxidant, anti-inflammatory, anti-obesity, anticancer activity, and can decrease the lipid profile (Bunea et al., 2008; Gutierrez et al., 2019; Roberts & Moreau, 2016). A more recent study by Perez Pinero et al., showed that 12 weeks of supplementation rich in spinach extract in elderly increased the muscle mass and boosted muscle strength (Pérez-Piñero et al., 2021). Similar observations were also made in young strength-experienced individuals (Isenmann et al., 2019).

The intake of quinoa and spinach results in the detection of ecdysterone and its metabolites in urine. Related to doping control studies, it may lead to positive findings even if athletes do not consume ecdysterone as a performance-enhancing agent. However, based on their nutritional values and benefits, the consumption of quinoa and spinach is inevitable. Therefore, it is necessary to investigate the urinary excretion of ecdysterone after the consumption of quinoa and/or spinach. In our previous study the intake of 18 to 19 mg of ecdysterone from spinach as sautéed or as smoothies resulted in the detection of ecdysterone in the urine with a maximum concentration range from 0.08 – 0.74 µg/mL (Yuliandra et al., 2023).

The aim of this study was first to test different brands of quinoa seeds for their ecdysterone concentration to then determine the ecdysterone concentration and its metabolites in urine after a single application of a high dietary ingestion of quinoa alone and in combination with spinach.

## 2. Material and Methods

### 2.1. Chemicals

Reference material of ecdysterone (2β,3β,14α,20β,22R,25-hexahydroxy-5β-cholest-7-en-6-one, purity > 95%) was obtained from Steraloids (Newport, RI, USA). Ponasterone (2β,3β,14α,20β,22R-pentahydroxy-5β-cholest-7-en-6-one) used as an internal standard (ISTD), was bought from Santa Cruz Biotechnology, Inc. (Heidelberg, Deutschland). Alpha-14-deoxy-ecdysterone and alpha-14-deoxy-poststerone were purchased from Extrasynthese (Genay CEDEX, France). Stock solutions of the analytes in methanol were prepared at a concentration of 1 mg/mL and stored at −20° C until further use.

### 2.2. Food Analysis

Prior to the use and ingestion of quinoa, a total of five different commercial quinoa seeds were tested for their ecdysterone concentration. The following brands were analyzed for this purpose:

> Quinoa 1: Farm Streit Köniz, m = 500 g
>
> Quinoa 2: Drugstore market (dm) Organic Quinoa Tricolore, m = 500 g
>
> Quinoa 3: Drugstore market (dm) Organic Quinoa normal, m = 500 g
>
> Quinoa 4: Drugstore market (dm) Organic Quinoa puffed, m = 500 g
>
> Quinoa 5: Alnatura, organic quality, m=500 g

Aliquots of each cooked quinoa portion (corresponding to 2 g of raw quinoa) were supplemented with a mixture of ethanol:water (80:20, v:v) to a final volume of 35 mL. The mixture was homogenized with Ultra-Turrax T 25 basic (IKA^®^ WERKE, Germany) for 3 min at 19.000 – 24.000 min^-1^ and centrifuged at 3011 RCF for 10 min. The supernatant was collected, and the residue was re-extracted under the same conditions two more times to ensure the maximum extraction of ecdysterone. The combined extracts were concentrated in vacuum at 60° C and afterwards suspended in 50 mL of water. Sequential extractions were performed with 3×50 mL hexane, ethyl acetate, and butanol as the organic phase. Each organic extract was then separately concentrated in vacuum at 60° C. The dry extract was reconstituted with 100 mL of methanol, diluted (1:50, *v:v*) with methanol, and filtered through a particle filter (0.2 μm syringe filter). Afterwards, 80 μL of the samples were spiked with 20 μL ISTD ponasterone (working solution 1000 ng/mL) and transferred to autosampler vials. For each sample, three replicates were prepared and analyzed.

The extraction and quantitation of ecdysterone from sautéed spinach were performed according to a previous study (Yuliandra et al., 2023). The same source of spinach was used in both studies.

### 2.3. Administration Studies

#### 2.3.1. Study Design

After analyzing the ecdysterone concentrations of the different quinoa varieties, a total of eight healthy very well-trained subjects (four females and four males) participated in the study. The participants were 27 (±3) years old, 174 (±9) cm tall, and weighed 76 (±15) kg. In the first application, the eight participants received only cooked quinoa. The sample yielding the highest ecdysterone concentration was used for administration. In the second intervention, participants consumed a combination of cooked quinoa and sautéed spinach leaves. There was a washout period of at least one week between the two applications.

All participants were instructed not to consume ecdysterone-containing foods before and during the trials. Blank urine was collected before both administrations. After administration, urine samples were collected until day 3 (72 h). Time of sample collection and urine volume were recorded. Aliquots of urine samples were stored frozen at −18 °C until analysis.

The study was approved by the local ethics committee of the German Sport University Cologne (No. 152/2020) and carried out in accordance to the provisions of the Declaration of Helsinki. All participants gave their written consent to participate in the study before the first application.

#### 2.3.2. Food Preparation

Quinoa (Alnatura, Darmstadt, Germany, lot: 13552) and frozen spinach (Globus, Cologne, Germany, lot: L21263-LN04) were purchased from the German local market. On the day of administration, 150 grams of raw quinoa was combined with twice the amount of water, then simmered in low heat for about 20 – 25 minutes. The sauteed spinach was prepared according to a previous study (Yuliandra et al., 2023). Subjects received portions of 150 g of raw quinoa at each meal which equals to an average of 368.8 (61) g of cooked quinoa. During the second intervention, they additionally received one packet (1 kg according to the manufacturer) of spinach, equivalent to 809 (115) g of sautéed spinach. Before the ingestion, each portion was weighed and recorded.

#### 2.3.3. Preparation of Calibration Standards and Quality Control Samples

A working solution of ecdysterone was prepared as in previous studies (Yuliandra et al., 2023). Dilutions of the stock solution with methanol were prepared to produce 5, 10, 25, 50, 250, 500, 1250, and 2500 ng/mL concentrations. The ISTD working solution was prepared by diluting the ponasterone stock solution with methanol to achieve a concentration of 1000 ng/mL. Calibration media were prepared by adding 20 μL ponasterone (ISTD) and 20 µL of the respective ecdysterone working solutions to 60 µL methanol. The following final concentrations of ecdysterone for calibration were used: 1, 2, 5, 10, 50, 100, 250, and 500 ng/mL. All calibration points were produced in duplicates.

### 2.4. Urine Analysis

#### 2.4.1. Preparation of Calibration Standards and Quality Control Samples

The detailed procedure for the preparation of matrix-matched calibrations was described in previous study (Ambrosio et al., 2021). Blank urine samples (200 μL) were spiked with 10 µL of each ecdysterone, 14-deoxy-ecdysterone, 14-deoxy-poststerone, and ponasterone (ISTD, 10 µg/mL) and diluted with 760 µL methanol: water (MeOH: H_2_O, 10:90) resulting in final concentrations of matrix-matched calibrants from 1 to 5000 ng/mL. For ecdysterone, the calibration range was constructed from 2.5 – 2500 ng/mL, while for the metabolites, 14-deoxy-ecdysterone and 14-deoxy-poststrerone, the calibration range was constructed from 2.5 – 500 ng/mL. All calibration points were prepared in duplicate.

#### 2.4.2. Sample Preparation

The urine samples were prepared as reported previously (Ambrosio et al., 2021). In brief, urine samples (200 μL) were spiked with 10 μL of ISTD ponasterone (10 µg/mL) and diluted to 1 mL with MeOH: H_2_O (10:90 *v/v*). After vortex-mix and centrifugation at 9677 RCF for 8 minutes, the supernatants were transferred to vials. Aliquots of 5 μL of samples were injected into the LC-MS system for analysis.

### 2.5. Instrumentation

The instrumental analyses were performed by ultrahigh performance liquid chromatography-tandem mass spectrometry (UHPLC-MS/MS) system consisting of an Agilent 1290 Infinity II UHPLC coupled to an Agilent 6495 triple quadrupole tandem MS system (Agilent Technologies GmbH, Waldbronn, Germany), utilizing an Agilent Jet Stream electrospray ionization (ESI) source and Ion Funnel. The same LC-MS/MS method was used for the quantitation of ecdysterone in food and ecdysterone and its metabolites in urine samples, with a different LC column and flow rate of the mobile phase.

Chromatographic separation was achieved using an Agilent Eclipse Plus C18 column (2.1 mm × 100 mm, particle size 1.8 µm) in food analysis, and an Agilent Eclipse Plus C18 column (2.1 mm × 50 mm, particle size 1.8 µm) in urine analysis.

The mobile phase comprised of formic acid in water (H_2_O:FoOH 99.9:0.1 *v:v*, eluent A) and formic acid in acetonitrile (ACN:FoOH 99.9:0.1 *v/v*, eluent B). For food analysis, the gradient program started at 10% eluent B for 2 min, linearly increased to 90% in 4 min, 1 min hold, followed by 0.50 min back to 10% eluent B for re-equilibration. For urine analysis, the gradient program started at 12% of eluent B and linearly increased to 40% in 4 min, then to 98% in 1.20 min, 0.30 min hold, followed by 0.20 min re-equilibration at 12% of eluent B. For food analysis, the flow rate was 0.5 mL/min and the total run time was 7.5 min plus 2.5 min after each run for column equilibration. Aberrantly, for urine analysis a flow rate of 0.45 mL/min was used resulting in a total run time of 5.7 min plus 2 min for column equilibration. The sample injection volume was 5 µL and the temperature of the autosampler was maintained at 5° C.

The triple-quadrupole mass spectrometer (QqQ) was operated using positive electrospray ionization (ESI+). A capillary voltage of 3,500 V, a nozzle voltage of 300 V, a drying gas flow of 15 L/min (nitrogen) at 150° C, a sheath gas flow of 12 L/min (nitrogen) at 375° C and a nebulizer pressure of 25 psi (nitrogen) were used. The protonated molecular ion [M+H] ^+^ for ecdysterone was detected at *m/z* 481, for 14-deoxy-ecdysterone and ponasterone at *m/z* 465 (isomers) and at *m/z* 347 for 14-deoxy-poststerone. For MS/MS nitrogen was used as collision gas. Data acquisition in multiple reaction monitoring (MRM) mode was used for the entire study.

Detailed MS parameters for MRM transitions are summarized in supplement 1: Table S1 for food analysis and in Table S2 for urine analysis. Method performance was characterized as reported earlier and complemented as reported in supplement 2 (Figure S1 and Table S3).

### 2.6. Software

MassHunter software version 10 from Agilent was used for data acquisition. MassHunter Quant software version 10 from Agilent was used for data processing. Microsoft Excel 365 was used for data analysis and data visualization. OriginPro, version 2019b (OriginLab Corporation, Northampton, MA, USA) was used for statistical analysis and data visualization.

### 2.7. Evaluation of Urinary Data

The parameters of urinary excretion kinetics of ecdysterone, 14-deoxy-ecdysterone, and 14-deoxy-poststerone were evaluated as previously described (Ambrosio et al., 2021). In brief, the concentration of the three analytes were corrected with the dilution factor (1:5). The excretion rate (E_rate_), the half-life, and the cumulative amount of three analytes were calculated as previously described in detail (Ambrosio et al., 2021). The recovered amounts of analytes in the urines is reported in percentage in comparison with the intake of ecdysterone in each subject.

### 2.8. Statistical Analysis

The anthropometric data of participants, the intake of food, and quantitation in quinoa and spinach were reported as mean (standard deviation, SD). The urinary parameters such as maximum concentration (C_max_), maximum excretion rate (E_rate-max_), cumulative amount, recovered amount (%), and half-life were reported as a range (min–max), as mean (SD), and/or as median (IQR, interquartile range). Samples with n=3 were assumed to be not normally distributed due to a low number of samples. Samples (n> 3) were tested with Shapiro-Wilk for normal distribution. The paired t-test was used to evaluate the statistical difference for normally distributed data. The nonparametric Wilcoxon signed-rank test was used to evaluate the statistical difference for non-normally distributed data. A p-value < 0.05 was considered significant.

## 3. Results

### 3.1. Ecdysterone Amount in Different Quinoa Varieties and Spinach

The analysis of the amount of ecdysterone from the five quinoa varieties showed that the highest ecdysterone concentration is obtained in the quinoa from Alnatura. The ecdysterone concentration from the sauteed spinach (FW) was 20.2 (0.3) µg/g. All ecdysterone concentrations are listed in supplement 3 Table S4.

Therefore, according to the intake of each subject (equal to 150 g raw quinoa), the participants consumed 55.3 mg of ecdysterone in the first study with only quinoa. In the second study, the intake of ecdysterone based on the sum of quinoa intake and spinach intake was 71.6 (2.3) mg.

### 3.2. Urinary Excretion Parameters of Ecdysterone and Its Metabolites

The concentration over time (A), the excretion rate against the midpoint time of sample collection (B), and the cumulative amount (C) after the intake of quinoa (upper) and the combined intake with spinach (lower) for ecdysterone, 14-deoxy-ecdysterone, and 14-deoxy-poststerone are illustrated in Figure 1, Figure 2, and Figure 3.

**Figure 1:**
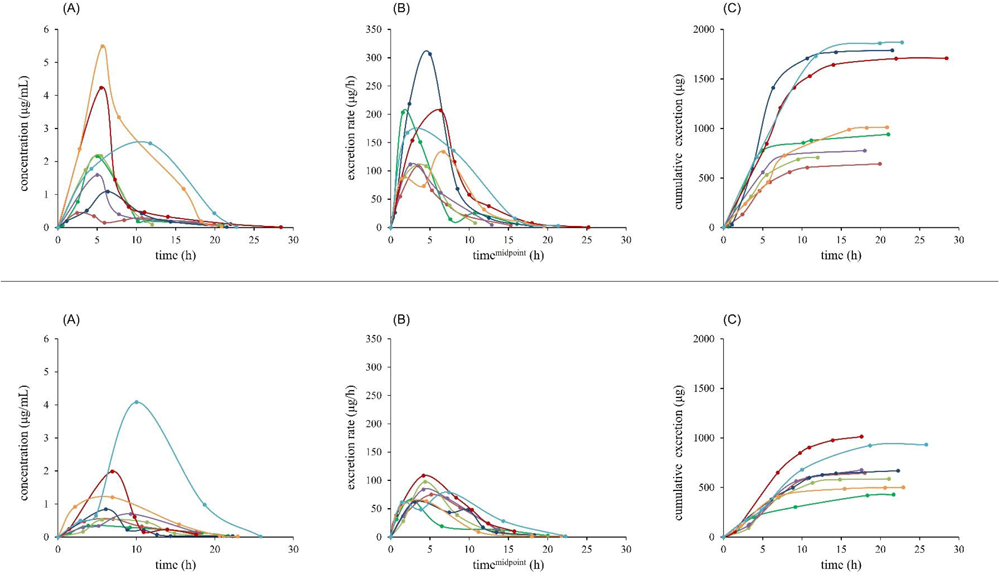
Urinary excretion profile of ecdysterone: concentration over time (A), excretion rate against midpoint time of sample collection (B), and cumulative amount (C) after the intake of cooked quinoa alone (upper) and the combination of cooked quinoa and sautéed spinach (bottom). Each color represents each subject.

**Figure 2:**
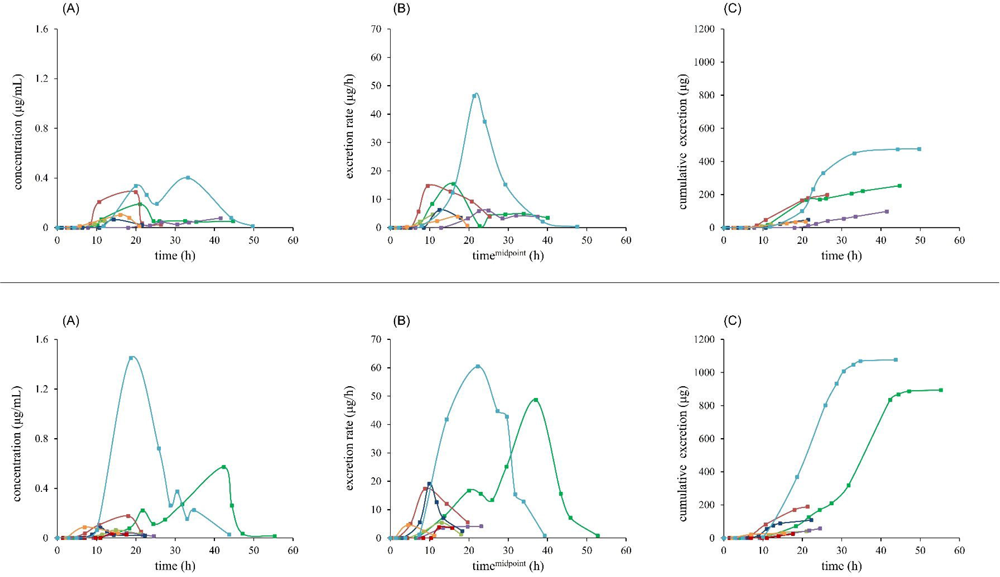
Urinary excretion profile of 14-deoxy-ecdysterone: concentration over time (A), excretion rate against midpoint time of sample collection (B), and cumulative amount (C) after the intake of cooked quinoa alone (upper) and the combination of cooked quinoa and sautéed spinach (bottom). Each color represents each subject.

**Figure 3:**
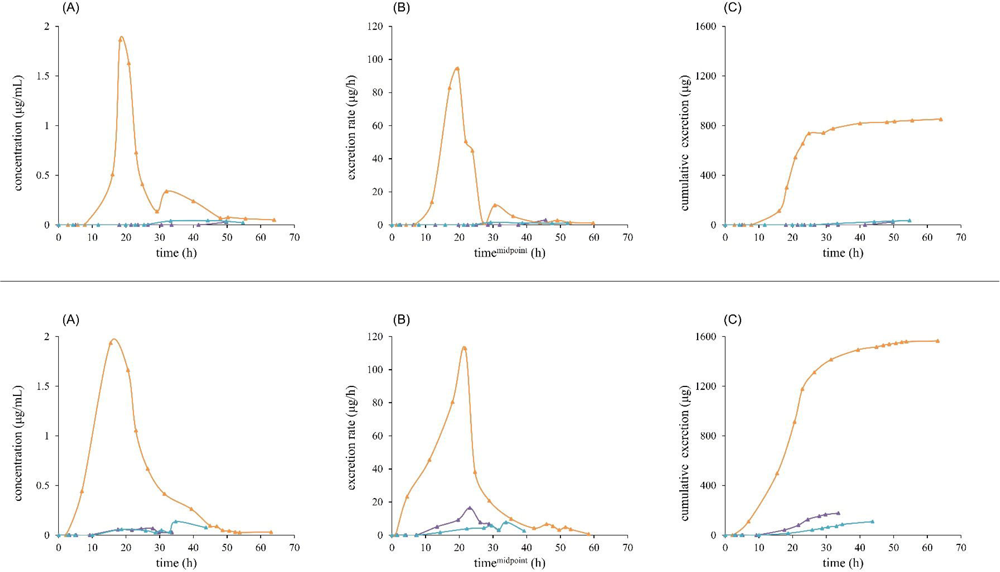
Urinary excretion profile of 14-deoxy-poststerone: concentration over time (A), excretion rate against midpoint time of sample collection (B), and cumulative amount (C) after the intake of cooked quinoa alone (upper) and the combination of cooked quinoa and sautéed spinach (bottom). Each color represents each subject.

Following the intake of cooked quinoa alone, ecdysterone was excreted in the urine and reached the maximum concentration in the range of 0.44 – 5.50 µg/mL achieved between 2.4 – 12 h. In the concomitant administration with spinach, the maximum concentration was achieved between 4 – 10 h with the concentration obtained in almost similar range (0.34 – 4.09 µg/mL). The maximum excretion rate of ecdysterone for quinoa intake was found in the range of 109 – 307 µg/h in the 1.5 – 7 h samples (midpoint time), while for spinach plus quinoa ingestion, the maximum excretion rate was lower (62 – 108 µg/h) and reached between 3 – 7 h midpoint time. The cumulative amount of ecdysterone was 642 – 1870 µg following quinoa intake, and 428 – 1013 µg after the intake of spinach plus quinoa.

The metabolite of ecdysterone, 14-deoxy-ecdysterone, achieved its maximum concentration in urine between 12 – 42 h and the concentration obtained was 0.05 – 0.4 µg/mL after quinoa administration. In the second study, the maximum concentration was obtained in the range of 0.02 – 1.45 µg/mL at urine sampling times between 7 – 42 h. The maximum excretion rate was in the range of 4 – 47 µg/h and obtained between 10 – 25 h midpoint sampling time for quinoa. In the second administration, it was at 4 – 61 µg/h and achieved in the 5 – 37 h midpoint time. The cumulative excreted amount of 14-deoxy-ecdysterone was 20 – 476 µg and 25 – 1077 µg following quinoa and combined administration of quinoa and spinach, respectively.

For the second metabolite, 14-deoxy-poststerone, the maximum concentration in both intakes was highly similar (0.03 – 1.87 µg/mL for quinoa intake and 0.07 – 1.94 µg/mL for the combination intake). The maximum concentration was detected between 18 – 50 h post-administration in the first intake and 15 – 35 h in the second administration. The maximum excretion rate was 1.5 – 95 µg/h at 20 – 46 h (midpoint sampling time) for quinoa and between 7.8 – 113 µg/h achieved at 22 – 34 h (midpoint sampling time) for combined intake. The cumulative amount of 14-deoxy-poststerone in urine was 25 – 853 µg and 110 – 1565 µg for the first and second consumption, respectively.

The boxplot of the individual recovered amount (%) in urine (A) and the total excreted amount (%) in urine after the first and second intake is displayed in Figure 4. The mean (SD) of total excreted amount (sum of ecdysterone and two metabolites) was 2.60 (1.09)% following the quinoa intake and 1.71 (0.86)% after the combined intake with spinach. The paired t-test revealed a significant difference (p<0.05) in the total excreted amount (%) in urine between the quinoa intake and the combined intake with spinach. The mean (SD) recovered amount (%) of ecdysterone, 14-deoxy-ecdysterone, and 14-deoxy-poststerone was 2.14 (0.94), 0.29 (0.30), 0.55 (0.86) for quinoa intake alone and 0.96 (0.31), 0.43 (0.62), 0.85 (1.12) for the combined intake of quinoa and spinach, respectively.

**Figure 4:**
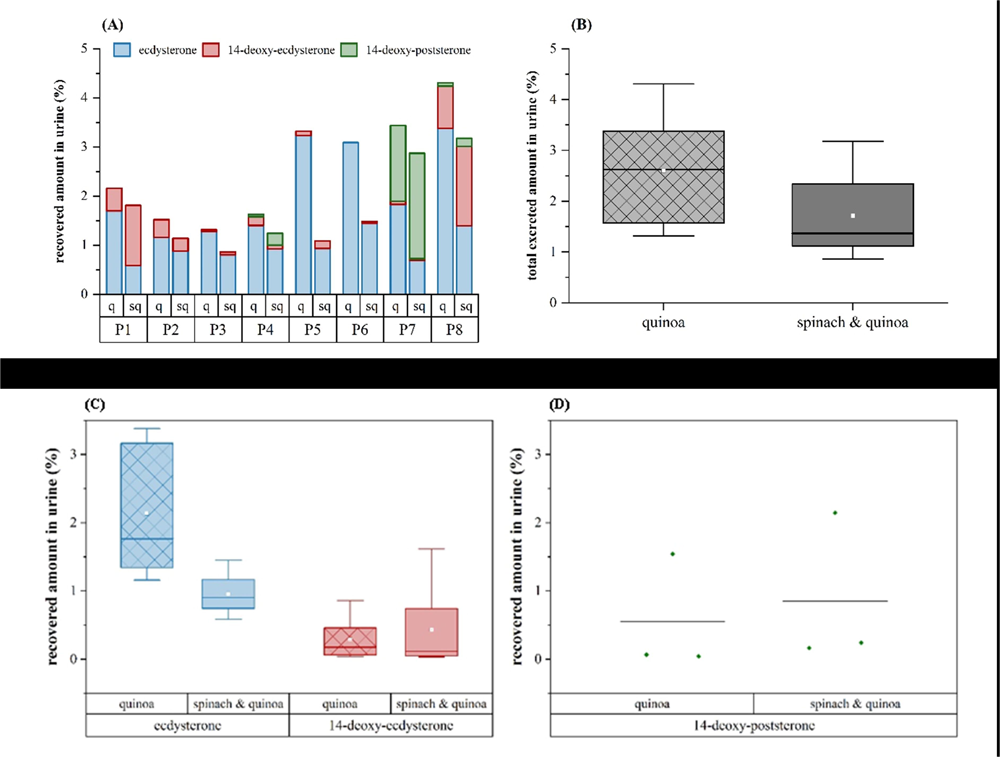
Box plots of recovered amount in urine, (A) individual recovered amounts (%), (B) total excreted amount (%) in urine, (C) ecdysterone and 14-deoxy-ecdysterone combined for all participants, (D) individual points for 14-deoxy-poststerone.

In most subjects, ecdysterone was excreted as major compound, while a few other participants excreted 14-deoxy-ecdysterone or 14-deoxy-poststerone as dominant compound indicating the interindividual variation between subjects.

The summary of urinary excretion parameters is presented in Table 1 for quinoa intake and in Table 2 for quinoa plus spinach ingestion.

**Table 1:** Quinoa ingestion: Summary of urinary elimination data of ecdysterone and its metabolites 14-deoxy-ecdysterone and 14-deoxy-poststerone.

|  | Quinoa |  |  |  |
| --- | --- | --- | --- | --- |
|  | n | Range | Mean (SD) | Median (IQR) |
| **Ecdysterone** | 8 |  |  |  |
| C<sub>max</sub> (µg/mL) |  | 0.44 – 5.50 | 2.47 (1.66) | 2.17 (2.05) |
| E<sub>rate-max</sub> (µg/h) |  | 109 – 307 | 169 (69) | 151 (95) |
| Cumulative amount (µg) |  | 642 – 1870 | 1181 (520) | 977 (1008) |
| recovered in urine (%) |  | 1.16 – 3.38 | 2.14 (0.94) | 1.77 (1.82) |
| **14-deoxy-ecdysterone** | 7 |  |  |  |
| C<sub>max</sub> (µg/mL) |  | 0.05 – 0.40 | 0.17 (0.13) | 0.1 (0.22) |
| E<sub>rate-max</sub> (µg/h) |  | 4 – 47 | 14. (15) | 6.26 (10.7) |
| Cumulative amount (µg) |  | 20 – 476 | 161 (164) | 98 (218) |
| recovered in urine (%) |  | 0.04 – 0.86 | 0.29 (0.30) | 0.18 (0.39) |
| **14-deoxy-poststerone** | 3 |  |  |  |
| C<sub>max</sub> (µg/mL) |  | 0.03 – 1.87 | 0.65 (1.06) | 0.04 (1.84) |
| E<sub>rate-max</sub> (µg/h) |  | 1.5 – 95 | 33 (53.4) | 2.97 (93.1) |
| Cumulative amount (µg) |  | 25 – 853 | 305 (475) | 36.5 (828) |
| recovered in urine (%) |  | 0.04 – 1.52 | 0.55 (0.86) | 0.07 (1.48) |

n= number of participants who excreted the compound in urine

**Table 2:** Quinoa in combination with spinach ingestion: Summary of urinary elimination data of ecdysterone and its metabolites 14-deoxy-ecdysterone and 14-deoxy-poststerone.

|  | Quinoa + Spinach |  |  |  |
| --- | --- | --- | --- | --- |
|  | n | Range | Mean (SD) | Median (IQR) |
| **Ecdysterone** | 8 |  |  |  |
| C<sub>max</sub> (µg/mL) |  | 0.34 – 4.09 | 1.28 (1.25) | 0.77 (1.06) |
| E<sub>rate-max</sub> (µg/h) |  | 62 – 108 | 79.4 (16.7) | 77 (26.1) |
| Cumulative amount (µg) |  | 428 – 1013 | 682 (200) | 657 (261) |
| recovered in urine (%) |  | 0.59 – 1.45 | 0.96 (0.31) | 0.9 (0.42) |
| **14-deoxy-ecdysterone** | 8 |  |  |  |
| C<sub>max</sub> (µg/mL) |  | 0.02 – 1.45 | 0.31 (0.49) | 0.09 (0.33) |
| E<sub>rate-max</sub> (µg/h) |  | 4 – 61 | 20.5 (22.1) | 11.4 (29.6) |
| Cumulative amount (µg) |  | 25 – 1077 | 303 (428) | 83 (507) |
| recovered in urine (%) |  | 0.04 – 1.6 | 0.43 (0.62) | 0.12 (0.70) |
| **14-deoxy-poststerone** | 3 |  |  |  |
| C<sub>max</sub> (µg/mL) |  | 0.07 – 1.94 | 0.71 (1.06) | 0.14 (1.87) |
| E<sub>rate-max</sub> (µg/h) |  | 7.8 – 113 | 45.7 (58.3) | 16.5 (105.1) |
| Cumulative amount (µg) |  | 110 – 1565 | 618 (822) | 177 (1456) |
| recovered in urine (%) |  | 0.17 – 2.15 | 0.85 (1.12) | 0.24 (1.98) |

n= number of participants who excreted the compound in urine

## 4. Discussion

Quinoa and spinach are two edible plants rich in ecdysterone in varying amounts (Bajkacz et al., 2020; Bakrim et al., 2008; Chen & Feng, 2021; Gorelick et al., 2020; Graf et al., 2014; Graf et al., 2016; Grebenok & Adler, 1991; Grucza et al., 2021; Kumpun et al., 2011). Since its inclusion in the WADA monitoring program (World Anti-Doping Agency, 2020), the urinary excretion of ecdysterone from food became a topic of interest. The present study aimed to investigate concentration of different quinoa varieties and the urinary excretion of ecdysterone and its metabolites in humans following the intake of quinoa alone and a combination intake with spinach. In this study, the intake of ecdysterone from quinoa was 55.3 mg, while the intake for the combination study with spinach was 71.6 (2.3) mg.

Ecdysterone was excreted and quantified in the urine of all participants after both applications. The maximum concentration after ingestion of quinoa only was slightly higher than in the combination with spinach, although the combined intake contained more ecdysterone. However, no significant differences were observed between the two study arms. Similar to the maximum concentration of ecdysterone, the maximum excretion rate and the cumulative amount of ecdysterone were two to three times higher in the first intake than in the combination intake. Most likely due to the high interindividual variety of urinary flow (which strongly influences urinary concentrations) statistical significance was achieved. On the contrary, the maximum excretion rate and the cumulative amount of ecdysterone revealed significant differences between the first intake and the second intake (p<0.05).

Ecdysterone is excreted almost completely from the urine within 24 hours by all six participants in both applications. In two participants ecdysterone was still detectable after 26 to 28 hours after one application. The half-life of ecdysterone for both intakes ranged between 2 and 5 hours, which is consistent with previous studies (Ambrosio et al., 2021; Dioh et al., 2023; Yuliandra et al., 2023). The result also indicates that the parent compound is rapidly excreted from the body.

Comparing the total amounts of ecdysterone excreted with the dose ingested, only about half the amount of ecdysterone was found after the combination with spinach despite the higher total amount of ecdysterone. Amounts of 1.16 – 3.38% were recovered when quinoa alone was consumed and 0.59 – 1.45% when combined with spinach. Potential reasons for low recoveries in general might be related to the food matrix, the volume of the food matrix, and the nature of the food matrix.^48-50^ The combination of quinoa and spinach has an even more complex food matrix, which might explain the lower absorption after the second meal. However, further investigations about the mechanism of bioaccessibility of ecdysterone from food are needed to allow a better understanding of the absorption processes.

Slightly higher urinary recoveries after quinoa ingestion were observed in a pilot project of our research group (supplement 4). Six healthy individuals (three females, three males) consumed 119-184 g of quinoa porridge (Quinoa 3) corresponding to 19.29-29.68 mg of ecdysterone (details in supplement 4). Similar ecdysterone concentrations and excretion rates were observed in the six participants (supplement 4). The results of the pilot project still confirm that only a small amount of ecdysterone (1/12-1/20) is recovered in urine after ingestion of quinoa. Comparing the result with our previous study of a single oral administration of pure ecdysterone (Ambrosio et al., 2021), the ecdysterone recovery after quinoa or quinoa and spinach was only 1/8 and 1/19 of the pure ecdysterone intake (without food matrix), respectively. In our previous study, the maximum concentration of ecdysterone in urine was between 4.4 and 30 µg/mL after 50 mg of pure ecdysterone (Ambrosio et al., 2021). In contrast, after quinoa only intake the maximum concentration of ecdysterone was 0.44 – 5.50 µg/mL and in the combination intake with spinach was 0.34 – 4.09 µg/mL. Even though the dosage of ecdysterone from food in the combined application is higher than that of pure ecdysterone, the maximum concentration obtained from oral administration of pure ecdysterone is considerably higher than with food intake. However, the maximum concentrations of both interventions (quinoa alone and quinoa plus spinach) are considerably higher than for pure spinach application. Studies on the bioavailability and excretion of ecdysterone from sautéed spinach and smoothie demonstrated that the maximum concentration of ecdysterone ranged from 0.09 to 0.41 μg/mL after consumption of sautéed spinach and 0.08-0.74 μg/mL after consumption of smoothie. The total amount recovered in urine of the parent drug and metabolites is only 1.4 (1.0)% for both sauteed spinach and smoothie (Yuliandra et al., 2023). The result indicates that, in terms of actual bioavailability, both quinoa and spinach are poor sources of ecdysterone.

With regard to the metabolite 14-deoxy-ecdysterone, it was detected in the urine of all participants. However, in contrast to ecdysterone, the maximum concentration and excreted amount in urine (%) was slightly increased in the combined intake compared to quinoa intake alone, while the maximum excretion rate showed similar values between the two intakes. However, the differences were not significant in all three parameters. In terms of excretion time, this showed a maximum of 50 h (4 out of 8 within 24 h) when quinoa was applied alone and a maximum of 56 h (5 out of 8 within 24 h) when combined with spinach. But, only a small percentage of 14-deoxy-ecdysterone was excreted in the urine (< 1% in quinoa intake and < 2% in combination with spinach). In addition, as shown in Figure 2, high variations in the concentration, excretion rate, and cumulative amount of 14-deoxy-ecdysterone were observed between subjects. The interindividual variation might be related to the gut microbiota composition in each participant. The differences in diet, genetic information, environment, and health conditions will highly impact the composition of gut microbiota within an individual (Conlon & Bird, 2014).

Similar to our previous study only 3 out of 8 participants excreted the second metabolite, 14-deoxy-poststerone, both in quinoa intake and in combination intake with spinach (Yuliandra et al., 2023). All three participants were female and the same participants also excreted this metabolite in the spinach study (Yuliandra et al., 2023). The results suggest that there may be sex differences in the metabolism of ecdysterone and metabolite formation. However, further studies are needed to confirm this.

Similar to 14-deoxy-ecdysterone, 14-deoxy-poststerone showed a high interindividual variation of metabolism. One participant excreted a significantly higher concentration than the other two (Figure 3). In all three participants, the metabolite was still detected after two days. Similar to ecdysterone and 14-deoxy-ecdysterone, only a small amount of 14-deoxy-poststerone was excreted in the urine. No significant difference was observed with respect to the two treatments.

As a sum of the parent compound and two metabolites, the mean (SD) total amount excreted in urine was 2.60 (1.09)% after consumption of quinoa and 1.71 (0.86)% after combined consumption with spinach. A significant difference was observed between the two study arms, which was mainly due to the amount of ecdysterone excreted in the urine. As a comparison, the mean (SD) total excreted amount in urine (as unchanged and metabolites) after oral intake of pure ecdysterone was 21.1 (13.3)% (Ambrosio et al., 2021), and after sautéed spinach and smoothie intake was 1.4 (1.0)% (Yuliandra et al., 2023). The intake of ecdysterone either from quinoa alone or in combination with spinach revealed that only a small proportion of ecdysterone and its metabolites were recovered in urine. This underlines that urinary recoveries of ecdysterone and its metabolites are generally low, but even lower if administered as food ingredient.

## Acknowledgements

The authors thank Peter Broekmann, Steffen Loke, Sarah Valder, and Bernhard Wüst for inspiring discussions and technical assistance. All participants of the ingestion trials are acknowledged for their contribution to the study.

## Author Contributions

Conceptualization, M.K.P.; methodology, T.Y., K.T., M.B., and E.I.; formal analysis, T.Y., K.T. and M.K.P.; investigation, T.Y. and K.T.; resources, P.D., F.B., and M.K.P.; writing— original draft preparation, E.I., T.Y., and K.T.; writing—review and editing, M.K.P., F.B., and X.T.; visualization, T.Y. and K.T.; supervision, M.K.P., and F.B.; project administration, M.K.P.; funding acquisition, M.K.P., F.B., P.D., and X.T. All authors agree with submission and publication.

## Data Availability Statement

Raw data are stored by the authors.

