## supplements for "Quinoa as Functional Food? Urinary elimination of ecdysterone after consumption of quinoa alone and in combination with spinach"

### Supplement 1: Analytical Method Details – MS/MS conditions

*Table S1: Parameters for MRM transitions for ecdysterone and ISTD in food analysis.*

| Analytes | Retention Time (min) | Precursor Ion (*m/z*) | Product Ion (*m/z*) | Collision Energy | Polarity |
|---|---|---|---|---|---|
| **ecdysterone** | | | | | |
| quantifier | 3.71 | 481 | 371 | 12 | positive |
| qualifier | | 481 | 165 | 28 | positive |
| **ponasterone** | | | | | |
| quantifier | 4.43 | 465 | 109 | 32 | positive |
| qualifier | | 465 | 173 | 24 | positive |

*Table S2: Parameters for MRM transitions for ecdysterone, metabolites and ISTD in urine analysis.*

| Analytes | Retention Time (min) | Precursor Ion (*m/z*) | Product Ion (*m/z*) | Collision Energy | Polarity |
|---|---|---|---|---|---|
| **ecdysterone** | | | | | |
| quantifier | 1.92 | 481 | 445 | 12 | positive |
| qualifier 1 | | 481 | 371 | 12 | positive |
| qualifier 2 | | 481 | 165 | 28 | positive |
| **14-deoxy-ecdysterone** | | | | | |
| quantifier | 2.37 | 465 | 285 | 28 | positive |
| qualifier 1 | | 465 | 303 | 28 | positive |
| qualifier 2 | | 465 | 80.9 | 44 | positive |
| **14-deoxy-poststerone** | | | | | |
| quantifier | 2.97 | 347 | 329 | 25 | positive |
| qualifier 1 | | 347 | 311 | 25 | positive |
| qualifier 2 | | 347 | 173 | 20 | positive |
| **ponasterone** | | | | | |
| quantifier | 3.55 | 465 | 109 | 32 | positive |
| qualifier | | 465 | 173 | 24 | positive |

### Supplement 2: Analytical Method Performance Characterization

#### Selectivity

The analytical method was able to separate ecdysterone, 14-deoxy-ecdysterone, 14-deoxy-poststerone, and ponasterone. Furthermore, 14-deoxy-poststerone was chromatographically separated from the other three endogenous steroids (*viz.* isomers of 14-deoxy-poststerone): corticosterone (RT: 4.09 min), 11-deoxycortisol (RT: 4.26 min), 21-deoxycortisol (RT: 3.95 min), as well as two more unidentified endogenous isomers (RT: 3.06 min and 3.27 min), and thus quantified without interference.

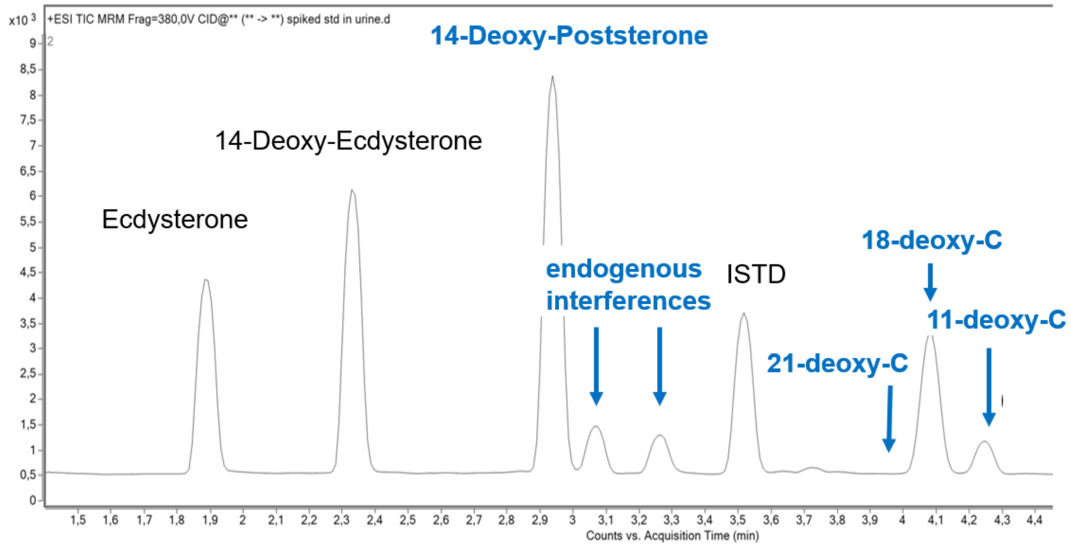


*Figure S1.1: Chromatogram of urine sample containing the analytes, internal standards and potentially interfering isomers displaying appropriate chromatographic separation of the endogenous isomeric compounds of 14-deoxy-poststerone, i.e. corticosterone (18-deoxy-C), 11-deoxycortisol (11-deoxy-C), 21-deoxycortisol (21-deoxy-C), as well as two more unidentified endogenous isomers (all marked in blue)*

#### Quality Control Samples

The quality control samples (QCs) were prepared independently at 3 different levels of ecdysterone concentration, *viz.* 5 ng/mL (low, LQC), 250 ng/mL (medium, MQC) and 500 ng/mL (high, HQC) for quinoa and spinach analysis.

For urine analysis the QCs were prepared independently at 4 different levels of concentration. The concentration of QCs for ecdysterone, 14-deoxyecdysterone, and 14-deoxypoststerone were 10/10/10 ng/mL (LQC), 250/100/100 ng/mL (MQC1), 1000/250/250 ng/mL (MQC2), and 2500/500/500 ng/mL (HQC), respectively.

#### Carryover

No carry-over observed in the blank urine following the injection of HQCs for all analytes.

#### Accuracy and Precision for Urine Sample Analysis

The accuracy and precision of LQC, MQC, and HQC during the run were all within acceptable criteria (accuracy: 85 – 115%, CV: <15%).

### Linearity of the Calibration Curves and LOD

*Table S3 Calibration Model, Calibration Range and LOD*

| Analyte | Calibration Model | Weighted | Calibration Range (ng/mL) | LOD (ng/mL) |
|---|---|---|---|---|
| Ecdysterone | Quadratic | 1/x | 2.5 – 2500 | 0.9 |
| 14-deoxy-ecdysterone | Linear | 1/x | 2.5 – 500 | 0.5 |
| 14-deoxy-poststerone | Linear | 1/x | 2.5 – 500 | 0.7 |

### Supplement 3: Analytical Method Performance Characterization

*Table S4: Ecdysterone content (µg/ g) from quinoa (DW) and sautéed spinach (FW) (n=3).*

| Type of Food | Mean (SD) [µg/g] |
|---|---|
| Quinoa 1#: Farm Streit Köniz | 122.33 |
| Quinoa 2#: DM Organic Quinoa Tricolore | 77.68 |
| Quinoa 3#: DM Organic Quinoa normal | 161.55 |
| Quinoa 4#: DM Organic Quinoa puffed | 72.43 |
| Quinoa 5#: Alnatura, organic quality | 369 (14)§ |
| Sautéed spinach | 20.2 (0.3)§ |

#calculated based on the raw quinoa (DW), §mean value (SD) of all consumed samples.

### Supplement 4: Pilot project quinoa application

#### Food preparation

At the beginning of the experiment, six subjects (three females, three males, M1, M2, M3, F1, F2, F3) consumed quinoa porridge up to high satiation for lunch. For food preparation, quinoa grains (Quinoa 3) were cooked with milk (500 g quinoa grains per 1 L of milk).

#### Ingestion and ecdysterone content

Based on the size of quinoa consumption and the concentration of ecdysterone in quinoa, the ingested amount of ecdysterone was calculated. The determination of the ecdysterone concentration was performed as described in the main manuscript. It was determined as 161.55 µg/g of raw quinoa grains. Table S1 lists the corresponding data for each individual.

*Table S5: Individual sizes of consumed quinoa portions in pilot project and corresponding ecdysterone amounts*

| Individual | Consumed amount of raw quinoa | Ingested amount of ecdysterone |
|---|---|---|
| M1 | 148.8 g | 24.04 mg |
| M2 | 167.4 g | 27.04 mg |
| M3 | 183.7 g | 29.68 mg |
| F1 | 140.6 g | 22.71 mg |
| F2 | 119.4 g | 19.29 mg |
| F3 | 135.4 g | 21.87 mg |

#### Results

In the post administration urines, concentrations of ecdysterone and 14-deoxyecdysterone determined by LC-MS/MS. The urinary concentrations against post administration time are displayed for each individual in Figure S2.2. As urinary flow has a direct impact on the concentration, excretion rates were determined to ease interpretation of the kinetics (Figure S2.3). Similar curves for the excretion rates are found for all subjects, with maximum rates observed in the first post-administration sample. Excretion of ecdysterone as parent substance was almost complete after 24 h. The proportion of ecdysterone parent decreases with time after administration, while the proportion of excreted metabolite increases.

Cumulative excretion data for both analytes (parent compound and metabolite 14-deoxyecdysterone) are plotted separately in Figure S2.4. It illustrates that while ecdysterone parent excretion was found almost complete after 24 h, 14-deoxyecdysterone was still detectable at the end of the collection period in all participants. Relevant increases in cumulative amounts of 14-deoxyecdysterone excretion were still observed in four of the six volunteers after 24 h.

Figure S2.5 and Figure S2.6 report the total amount of ecdysterone and its metabolite 14-deoxyecdysterone. Mean values of total recovered ecdysterone (as parent compound or 14-deoxyecdysterone) were determined as 11.0% within 24 h after administration.

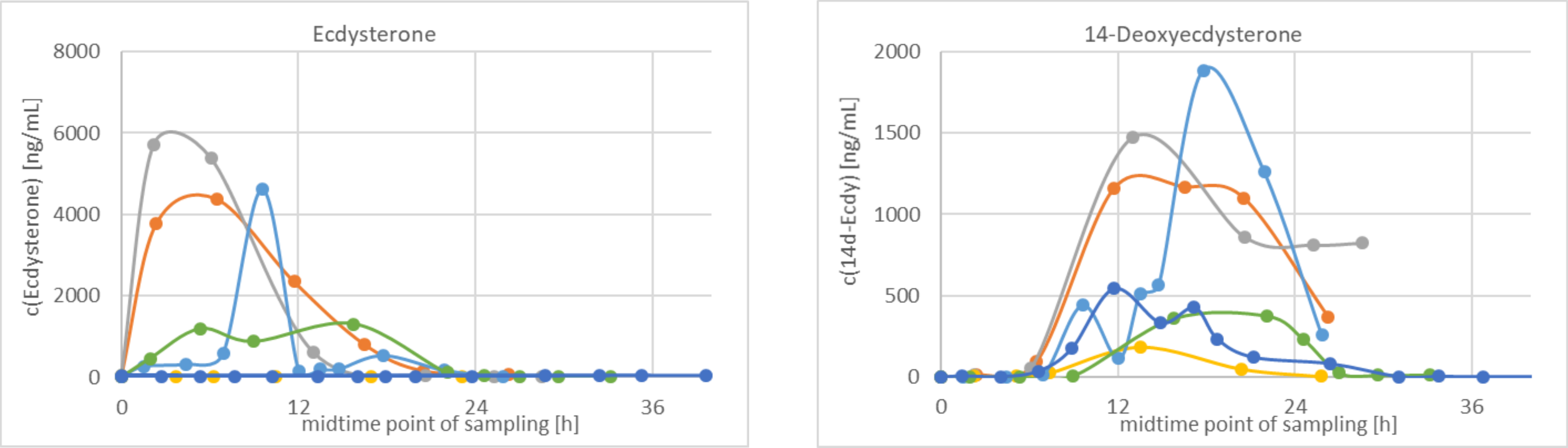


*Figure S2.2: Urinary profiles after quinoa porridge: concentration [ng/mL] of ecdysterone (left) and 14-deoxyecdysterone (right), M1 (red), M2 (grey), M3 (orange), F1 (light blue), F2 (green), F3 (dark blue)*

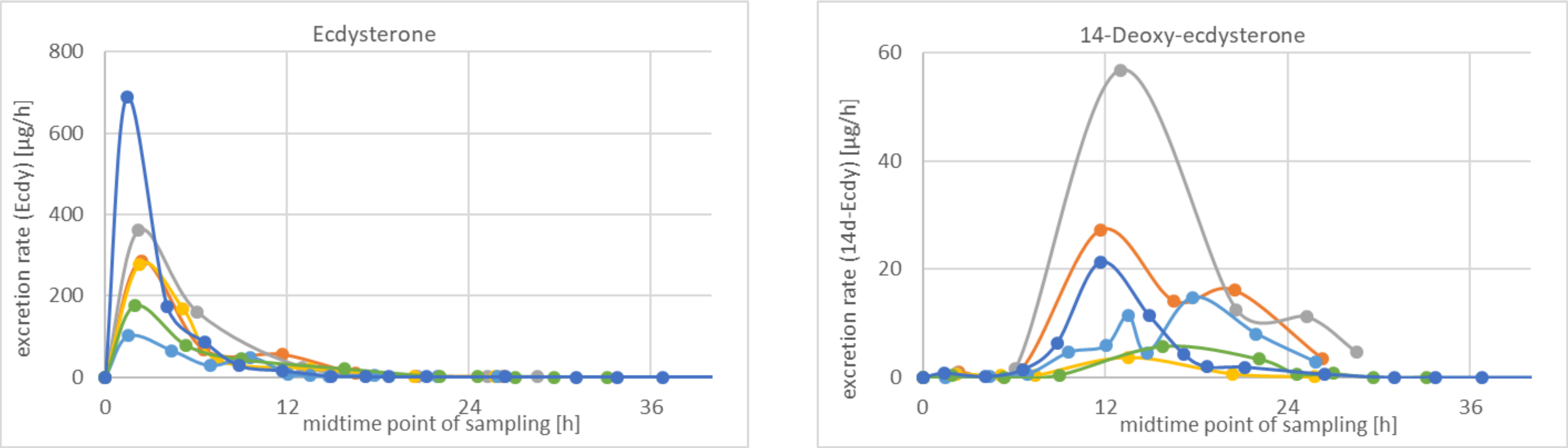


*Figure S2.3: Urinary excretion rate [µg/h] after quinoa porridge against midpoint time of sample collection: ecdysterone (upper) and 14-deoxyecdysterone (lower), M1 (red), M2 (grey), M3 (orange), F1 (light blue), F2 (green), F3 (dark blue)*

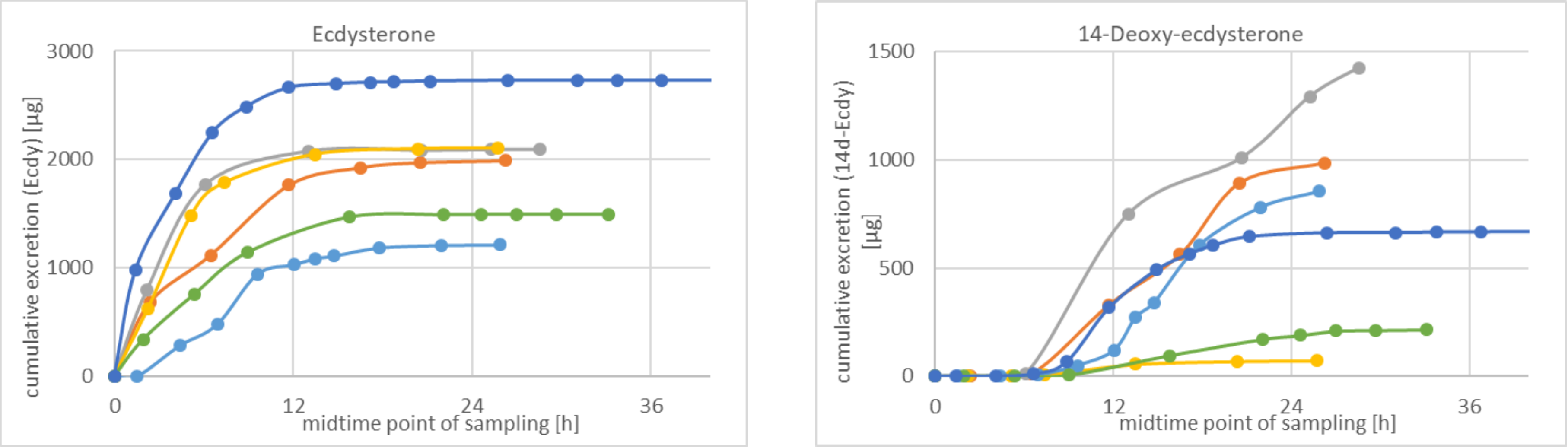


*Figure S2.4: Cumulative amount [µg] of urinary excreted ecdysterone (upper) and 14-deoxyecdysterone (lower) after quinoa porridge against midpoint time of sample collection, M1 (red), M2 (grey), M3 (orange), F1 (light blue), F2 (green), F3 (dark blue)*

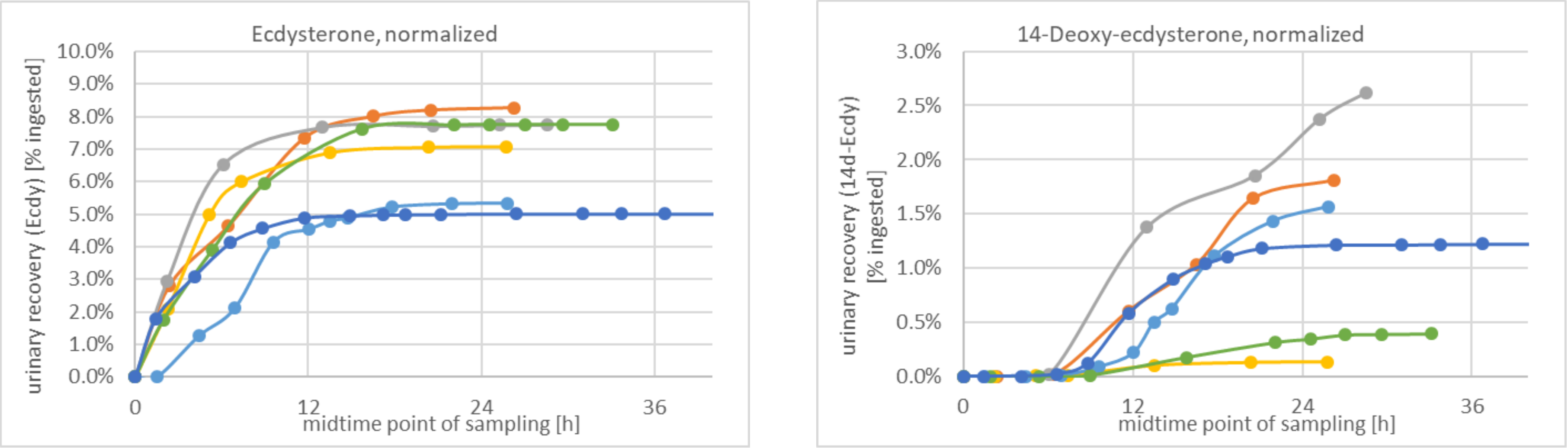


*Figure S2.5: Recovery [% of ingested ecdysterone] of urinary excreted ecdysterone (upper) and 14-deoxyecdysterone (lower) after quinoa porridge against midpoint time of sample collection, M1 (red), M2 (grey), M3 (orange), F1 (light blue), F2 (green), F3 (dark blue)*

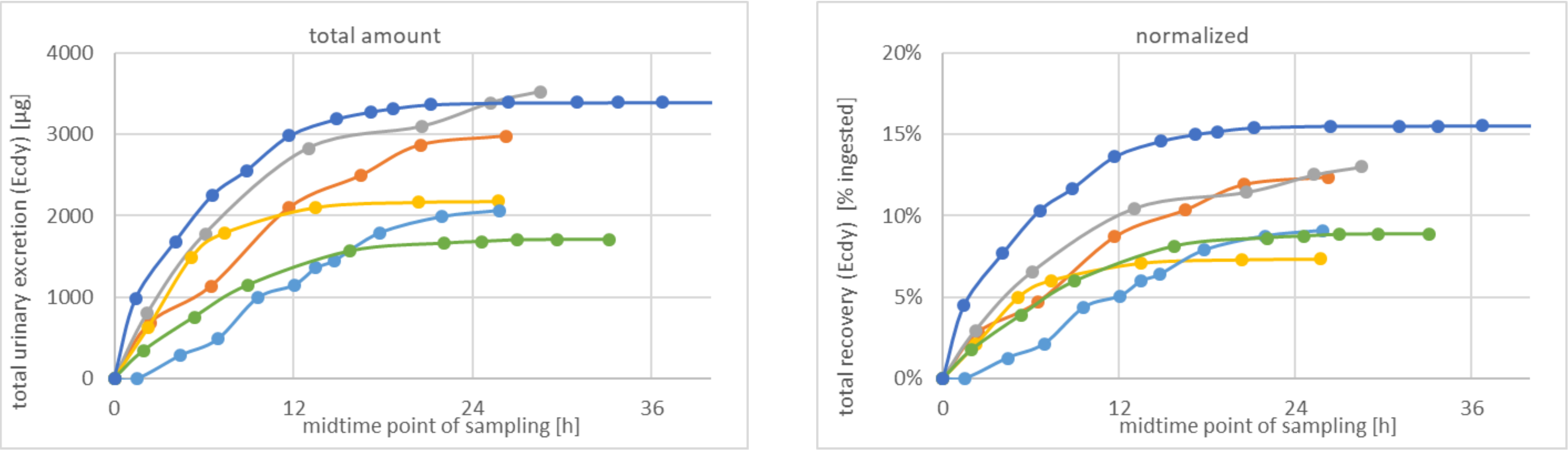


*Figure S2.6: Total cumulative urinary excretion of ecdysterone plus 14-deoxyecdysterone after quinoa porridge, amount [µg] (upper) and normalized [%] to ingested (lower), M1 (red), M2 (grey), M3 (orange), F1 (light blue), F2 (green), F3 (dark blue)*
